# Innate development of cognitive functions and motor programs by chemoaffinity

**DOI:** 10.64898/2026.01.30.702840

**Authors:** Friedrich Schuessler, Simone Ciceri, Henning Sprekeler

## Abstract

Humans and animals possess a rich repertoire of innate cognitive and behavioral abilities, yet the developmental mechanisms that establish the underlying neural circuitry remain unclear. Here, we show that a simple chemoaffinity-based connectivity rule, combined with stochastic cell-autonomous expression of cell adhesion molecules, can establish neural networks for a broad range of cognitive functions. We derive expression patterns that support the development of continuous attractor networks for path integration with toroidal grid-cell topology, networks with exponentially many discrete Hebbian assemblies, and circuits generating innate motor trajectories. Expression patterns can be combined in a modular way to encode recurrent networks for increasingly complex functions. Our results suggest that chemoaffinity may determine not only the anatomical organization of neural circuits, but also the structure of innate cognition and behavior.

---

Many complex behaviors with temporal dynamics are available shortly after birth, including spatial navigation and motor programs [1]. These behaviors are thought to rely on computations that unfold through recurrent neural interactions. Recurrent excitation in Hebbian assemblies completes patterns [2], recurrent inhibition can enforce decisions [3], networks with continuous attractors integrate evidence or velocity over time [4–7], and oscillatory or excitable dynamics generate rich motor patterns [8–10]. Signatures of such dynamics are observed before the onset of relevant sensory experience: head direction cells [11] and the grid cell attractor [12] are present before eye-opening and active navigation. Hence, both behavior and the recurrent dynamics that support it can be innate. How the underlying circuits are assembled is unclear [13].

One possibility is activity-dependent synaptic plasticity [14]. This need not require sensory experience: internally generated activity has been shown to play an important role in developmental circuit formation [15, 16]. In recurrent networks, however, synaptic plasticity is often unstable and requires fine-tuning even for relatively simple functions [17, 18]. Additionally, functional circuits can arise even in the absence of neural activity [19]. These observations suggest that genetically encoded, activity-independent programs contribute to the formation of recurrent circuits underlying innate behaviors.

In its early stages, neural circuit development relies on molecular cues. According to Roger Sperry’s idea of chemoaffinity, neurons form connections based on their genetically determined molecular identity [20]. Put simply, the formation and strength of synapses depend on whether a presynaptic “key” fits a corresponding postsynaptic “lock” (Fig. 1A). This idea was developed for retinotectal projections, where complementary molecular gradients align anatomical axes [Fig. 1B, [21]]. It has since undergone substantial refinement: keys and locks have been identified as sets of cell adhesion molecules acting in combinatorial patterns, and manipulating those patterns can rewire circuits in predictable ways [22, 23]. Most studies have considered chemoaffinity in the context of long-range, interareal connectivity [22]. Because keys and locks act locally, in a contact-based manner, the same mechanism could in principle also specify recurrent connections. Here we ask whether chemoaffinity can assemble recurrent networks for innate dynamical computations. We show that a simple molecular compatibility rule, combined with stochastic, cell-autonomous expression programs, is sufficient to establish continuous attractors for path integration, discrete neural assemblies, and circuits that generate sequences and motor trajectories. Expression programs can be combined modularly to form increasingly complex functions. Chemoaffinity may therefore not only provide a scaffold for wiring specificity, but also directly shape the dynamics of the recurrent networks that underlie innate cognition [24, 25] and motor behavior [1].

**Fig. 1:**
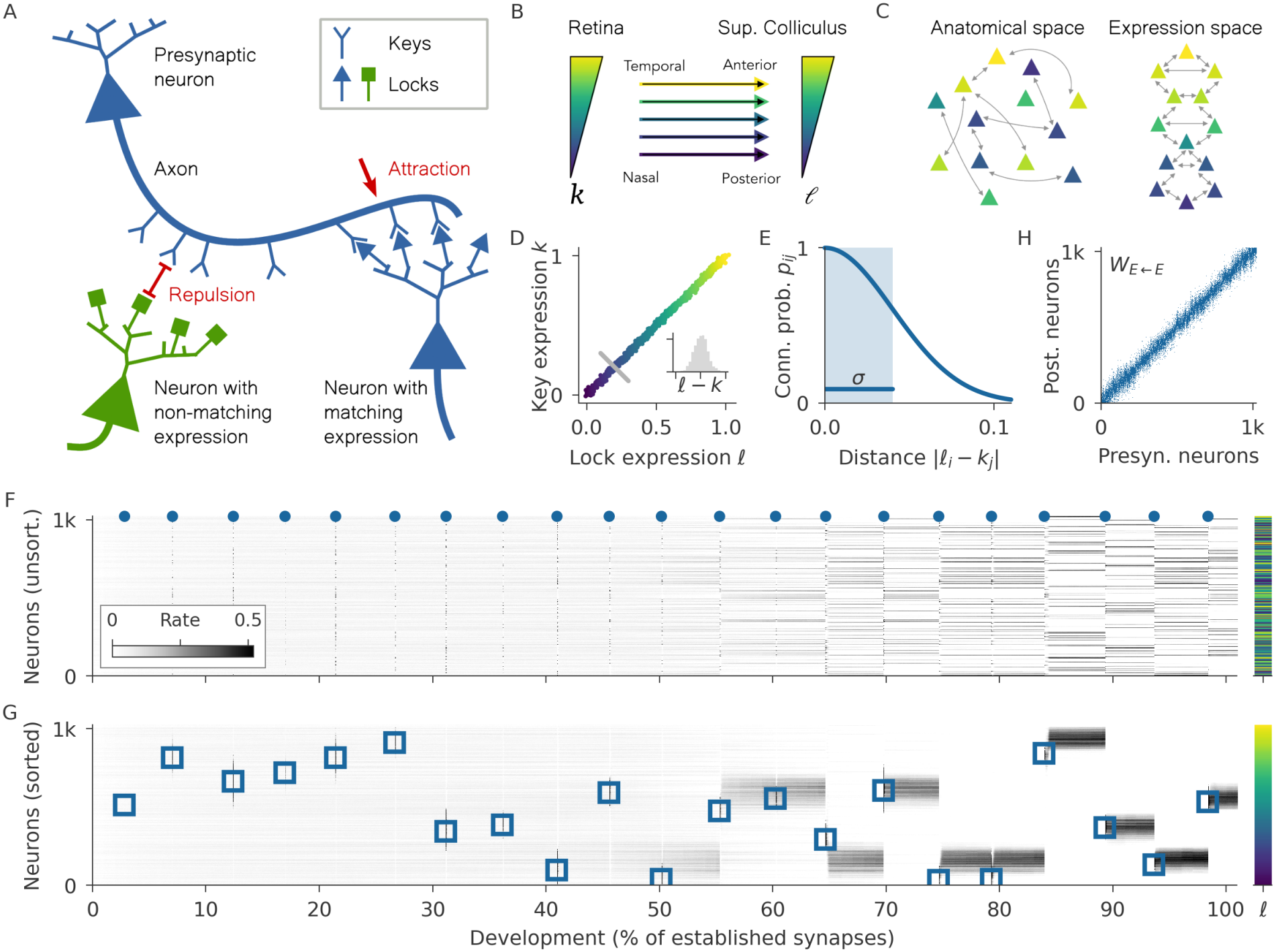
Innate development of a continuous attractor network by chemoaffinity. (**A**) Chemoaffinity: The axon of a presynaptic neuron (top) expresses a given key and is repelled by a neuron with nonmatching lock expression (bottom left), but attracted by a matching one (right). (**B**) Axon guidance, an instance of chemoaffinity, leads to a topographic mapping in the visual system. Axons from the retina connect to neurons in superior colliculus with matching concentrations of presynaptic receptors (acting as keys) and postsynaptic ligands (acting as locks) [26]. We used a phenomenological model, in which this matching requires similar expression levels of keys (*k*) and locks (*l*). In specific molecular systems, the matching condition may be different [27]. (**C**) Innate development of local recurrent connectivity by chemoaffinity. Expression levels may not correlate with neuronal position in anatomical space, but the resulting connectivity is local in expression space. (**D-G**) Innate development of a continuous attractor network. (**D**) Expression levels of locks and keys are correlated. Inset: Distribution of difference *l* − *k* between lock and key expression. (**E**) Connection probability as a function of distance in expression space. (**F**) Network activity over development, modeled by continuously adding synapses. Bar on the right indicates lock expression levels of each neuron. (**G**) Reordering the neurons by lock expression level reveals a local activity “bump”. In (**F, G**), the dynamics are repeatedly probed with brief input pulses (blue dots on top of **F**). These pulses are localized in expression space (blue squares in **G**). (**H**) Connectivity between excitatory neurons, ordered by lock expression levels, shows local excitation in expression space.

## Innate development of a continuous attractor network

Among the innate cognitive abilities known in mammals [25], the neural representation of space is arguably the best understood. In rodents, functional signatures of head direction cells [28] and grid cells [29] are found before eye opening and before the animals first leave their nest [11, 12, 30], suggesting that the underlying circuits are—at least initially—shaped by an innate developmental process. Both head direction and grid cells display collective activity patterns in the absence of sensory stimuli and are likely part of continuous attractor networks [31, 32]. How the highly structured recurrent connectivity required for these networks could be established innately is presently unknown [33].

To illustrate how chemoaffinity can establish a continuous attractor network, we first considered a “bump” attractor network as implied in place and head direction coding [4, 5, 34–37]. Bump attractor networks consist of neurons arranged in a line or ring that interact through local excitation and global inhibition. The resulting network dynamics establish a localized peak of persistent activity that can be shifted to different locations by external input. A challenge for these models is that anatomically nearby head direction and grid cells show weak correlations in their spatial tuning [29, 38], suggesting that the underlying network connectivity is not organized in anatomical space. Instead, we suggest that connectivity is structured in a molecular expression space by connecting neurons with matching expression levels of keys and locks (Fig. 1C).

We simulated an initially unconnected network of excitatory and inhibitory neurons. Each excitatory neuron expresses a stochastic amount *k* of a “key”, as well as a highly correlated amount *l* of the corresponding “lock” (Fig. 1D). These expression levels are cell-autonomous—that is, statistically independent across neurons—and uncorrelated with their anatomical position in the network. For conceptual simplicity, we use the term “keys” for presynaptic molecular labels, and “locks” for their postsynaptic counterpart. Over development, a chemoaffinity-based connectivity rule establishes new synaptic connections only between neurons for which the expression levels of the presynaptic key and the postsynaptic lock differ by less than a tolerance *σ* (Fig. 1E). To obtain global inhibition, the connections of the inhibitory neurons are formed randomly, without molecular specificity.

As more connections form, the initially unstructured network activity develops into a sparse activation pattern (Fig. 1F) with no obvious spatial structure. Ordering the neurons by their lock expression levels reveals the formation of a localized peak of activity (Fig. 1G). This bump reflects a continuous attractor, because it remains active in the absence of input and can be shifted to different locations in expression space by providing short input pulses to neurons with similar expression levels. Ordering neurons by expression levels also reveals local connectivity in expression space whose width is determined by the correlation between expression levels and the tolerance of connectivity [Fig. 1H, [27]]. As more connections form, the initially unstructured network activity develops into a sparse activation pattern (Fig. 1F) with no obvious spatial structure. Ordering the neurons by their lock expression levels reveals the formation of a localized peak of activity (Fig. 1G). This bump reflects a continuous attractor, because it remains active in the absence of input and can be shifted to different locations in expression space by providing short input pulses to neurons with similar expression levels. Ordering neurons by expression levels also reveals local connectivity in expression space whose width is determined by the correlation between expression levels and the tolerance of connectivity [Fig. 1H, [27]].

## An innate path-integration system with grid cells

Continuous attractor networks are the standard network model for path integration during spatial navigation [33]. In these models, recurrent excitation again generates a localized bump of activity, while directional biases in local connectivity allow velocity inputs to shift the position of the bump (Fig. 2A). Opposing biases in different subpopulations keep the bump stationary unless velocity inputs favor one population over another. When global inhibition is replaced by local inhibition, the network develops periodic activity patterns and the neurons become grid cells [6, 7].

**Fig. 2:**
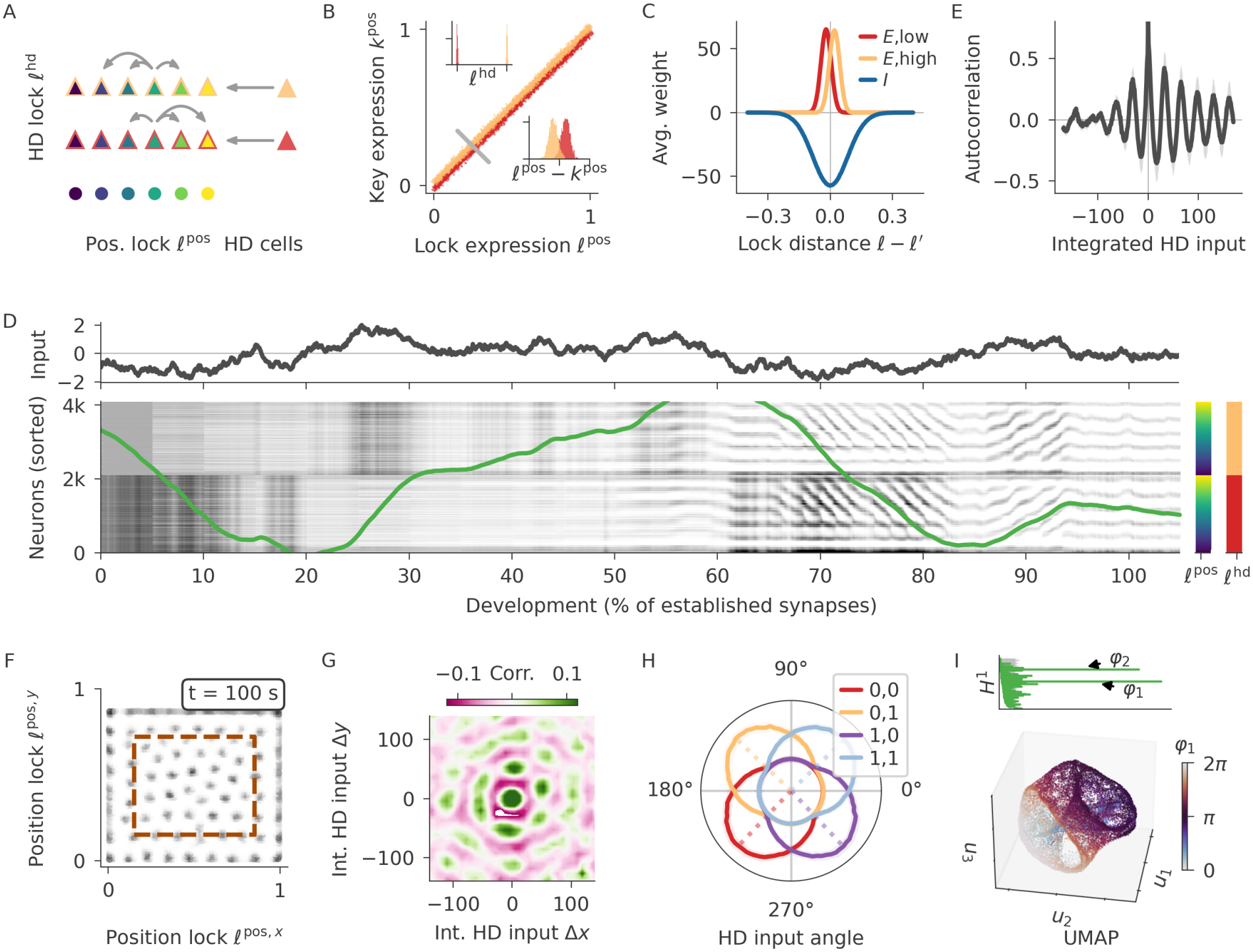
Innate development of a path-integration network. (**A-E**) 1-dimensional path-integration network. (**A**) Network structure: For excitatory neurons (triangles), the expression level of a head direction (HD) lock *l*^hd^ (border color) determines the input from HD neurons. Neurons connect to nearby neurons with a similar expression level of a positional lock *l*^pos^ (inner color). The HD lock biases the local connectivity to left (top) or right (middle). Inhibitory neurons (circles) also express the positional lock and key, but no head direction lock. (**B**) The expression of the positional key is correlated with the positional lock, but biased by the head direction lock. Insets: Distribution of the head direction lock (top) and the bias *l*^pos^ − *k*^pos^ (bottom). (**C**) Mean weight from excitatory (*E*) and inhibitory (*I*) neurons at a given distance in expression space. The molecular biases lead to biases in connectivity. Inhibitory connectivity is broader. (**D**) Top: HD input encoding velocity. Bottom: Network activity over development for excitatory neurons ordered by lock expression. At later stages of development, the activity is periodic and tracks the integrated input (green, scaled to match movement of bumps). (**E**) Autocorrelation of neural activity conditioned on integrated velocity reveals periodic activity pattern (mean and std. across excitatory neurons). (**F-I**) 2-dimensional path-integration network. (**F**) A snapshot of neuronal firing rates sorted according to the two position locks *l*^pos,*x*^, *l*^pos,*y*^. Neurons inside the dashed box show hexagonal patterns and are used for subsequent analyses. (**G**) Hexagonal autocorrelation of activity conditioned on integrated velocity along the two spatial dimensions. (**H**) Head direction tuning depending on the expression of the head direction locks *l*^hd,*x*^, *l*^hd,*y*^ (colors). Dotted lines indicate mean angle. (**I**) Toroidal topology in neural activity is revealed by two independent methods: Persistent cohomology (top) uncovers two rings with exceptionally long life times (x axis). Nonlinear dimensionality reduction (UMAP, bottom) visualizes a twisted torus [39]. Coloring by the cohomology angle *φ*_1_ shows consistency with the cohomology analysis.

Such a path-integration system can be formed by the same chemoaffinity rule by means of a low-dimensional and interpretable expression pattern of keys and locks. Consider a network of excitatory and inhibitory neurons (Fig. 2A), the connectivity of which develops based on two key-lock pairs: a “position” pair that organizes local connectivity and a “head direction” pair that organizes the velocity inputs. Each excitatory neuron expresses a random binary level of the head direction (HD) lock *l*^hd^ and thereby attracts inputs from a population of HD cells with the corresponding key expression (Fig. 2A). HD neurons with low and high key expression levels encode leftward and rightward movement along a linear track, respectively, once the animal leaves the nest. The excitatory neurons also express a “position” lock *l*^pos^ and the corresponding key *k*^pos^, both with continuous expression levels. The two are highly correlated, but the expression of the position key is slightly biased up or down by the HD lock *l*^hd^ (Fig. 2B). This molecular bias gives rise to the required biases in the local connectivity (Fig. 2C). Inhibitory cells only express the positional key and lock, generating the local inhibition required for the development of grid cells. The tolerance *σ* for deviations between key and lock expression is larger for inhibitory than for excitatory neurons, leading to effective Mexican hat-shaped interactions in the network.

We again simulated development by continuously adding synapses based on similarity in expression. Ordering the neurons by their expression levels reveals the emergence of a periodic activity pattern, the position of which starts shifting over time later during development (Fig. 2D). The position of this pattern is correlated with the integrated velocity encoded in the HD inputs, indicating that this system will perform path integration once the HD input represents the animal’s velocity (Fig. 2D, green). Individual neurons express periodic autocorrelation with respect to the integrated velocity, akin to grid cells in the entorhinal cortex [Fig. 2E, [29]].

The resemblance to the grid cell attractor in the entorhinal cortex [32] becomes even more pronounced when we extend the molecular system by two additional key-lock pairs that shape the encoding of a second spatial dimension. Ordering the neurons by the expression levels of the two position locks reveals a hexagonal activity pattern (Fig. 2F) that move in line with the integrated HD input (movie S1). Individual neurons display a hexagonal autocorrelation with respect to the integrated velocity (Fig. 2G), as well as head direction tuning (Fig. 2H), akin to conjunctive cells [40]. Finally, over the course of simulated development, the population activity shows the emergence of a toroidal topology [Fig. 2I, fig. S1, [39]], as observed in newborn rats [12]. In summary, a relatively low-dimensional chemoaffinity system can hard-code a fully functional path-integration system consisting of grid cells and head direction cells.

## Exponential expressivity: Discrete attractors, hierarchies & sequences

Given that a complex path-integration system can be formed by a relatively small number of key-lock pairs, we aimed to investigate how the complexity of the network scales with the number of molecules that encode its structure. To this end, we considered a system of *M* different key-lock pairs (Fig. 3A) with binary expression levels [23], because the complexity of the resulting networks can be readily quantified. The expression of keys and locks is again highly correlated (Fig. 3B).

**Fig. 3:**
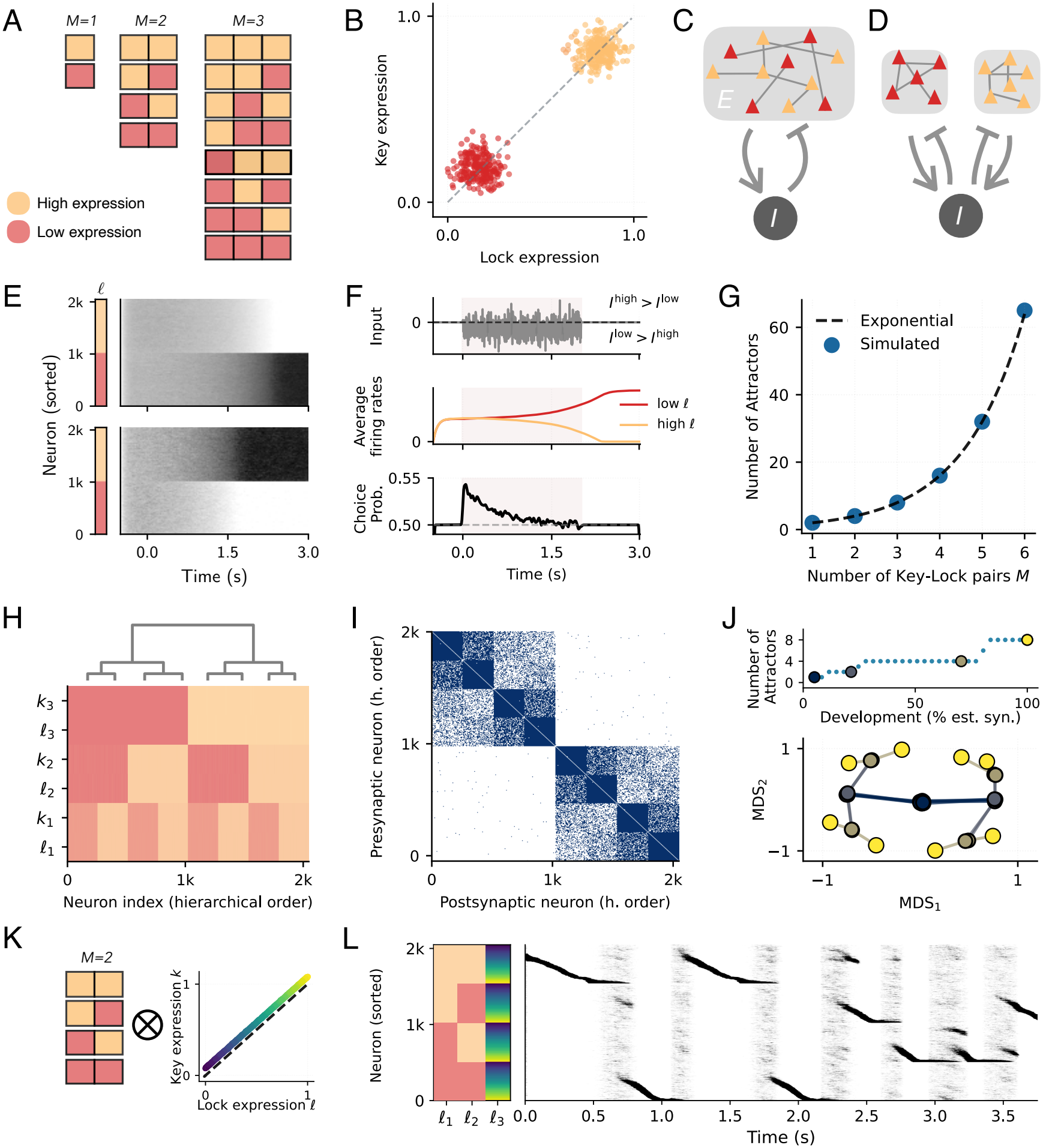
Discrete attractors, hierarchies and sequences. (**A**) Expression patterns for *M* = 1, 2, 3 different key-lock pairs. Colors indicate expression level. (**B**) lock and key expression levels are correlated and binary. (**C, D**) For one key-lock pair (*M* = 1), the recurrent network segregates into two assemblies. (**E**) The network activity displays two stable attractors when presented with different inputs (top vs. bottom). (**F**) When the two assemblies are both stimulated, the network integrates the difference of these two inputs. The choice probability indicates an integration over a time scale of a second. (**G**) The number of stable attractors scales exponentially with the number *M* of key-lock pairs. (**H-J**) Hierarchical networks. (**H**) Hierarchical clustering of the molecular expression levels (*M* = 3). (**I**) Connectivity matrix for neurons sorted according to the hierarchical clustering. (**J**) Top: During development, network activity initially falls into higher-order attractors, which subsequently split as more synaptic connections are established. Bottom: Multidimensional scaling (MDS) reveals a binary developmental tree. Colors indicate the developmental time points shown above. (**K**,**L**) Innate sequence model. (**K**) Combining expression patterns: two binary key-lock pairs (left), and one continuous key-lock pair with a bias between *k* and *l* (right; shift above diagonal). (**L**) Spontaneous activity of the network with neurons ordered by expression shows sequences. When a sequence reaches the end of the continuous expression profile within a subpopulation, the bump gradually decays and gives way to irregular activity until a new bump is formed in a random position and ignites a new sequence.

We expressed these guidance molecules in a network of excitatory neurons that were randomly connected to a population of inhibitory neurons and studied the structure and dynamics of the resulting network (Fig. 3C). For a single key-lock pair (*M* = 1), the excitatory population segregates into two unconnected assemblies with strong internal recurrent excitation (Fig. 3D). The resulting network dynamics reveal the presence of two stable attractors [Fig. 3E [2, 41]]. The network performs integration of a time-varying input (Fig. 3F), akin to network models for evidence integration during perceptual decision making [3].

When adding further key-lock pairs to the system, the number of stable attractors grows exponentially with the number of molecules (Fig. 3G). This result can be understood by interpreting the binary expression pattern of the guidance molecules as two M-item barcodes [42], one representing the expression pattern of keys and the other that of locks. Here, keys and locks are highly correlated, so these two barcodes are the same. Because there are 2^*M*^ different barcodes and only neurons with the same barcode forge connections, the network segregates into 2^*M*^ largely unconnected Hebbian assemblies. This suggests that the number of guidance molecules required for a given network architecture scales logarithmically with the complexity of the network [22].

The categorical specialization in these networks can be seen as a decision tree, where every key-lock pair induces a binary decision. Therefore, we reasoned that hierarchically clustered expression levels could generate networks whose activity reflects a hierarchy of cognitive categories [43]. Implementing such key-lock expression patterns (Fig. 3H) yields a network with hierarchically structured connectivity (Fig. 3I). The activity can settle into as many stable “low-level” attractors as there are molecular clusters (fig. S2). Multi-dimensional scaling of the network activity reveals that over development, these attractors emerge by splitting dynamics that reflect the structure of the hierarchy (Fig. 3J). Even after development is completed, different levels of the hierarchy can be accessed either by adding a readout organized by the same guidance molecules or by varying the noise level in the network dynamics (fig. S2).

The combinatorial expressivity of discrete key-lock pairs can be combined in a modular way with other molecular label systems to generate increasingly complex network dynamics. As an example, we simulated a network that combines discrete expression levels with a single continuously distributed key-lock pair that has a biased relation between keys and locks (Fig. 3K). After development, the network generates an exponentially large set of innate sequences (Fig. 3L), as in hippocampal preplay [44, 45].

## Innate motor programs

Given that chemoaffinity can generate temporal sequences, we reasoned that it may also enable the innate development of temporally structured motor programs. To explore this idea, we again considered a network with random, continuous expression levels of a lock *l*. If the expression of the associated key were the same as that of the lock, the network would display a stable bump of activity at an arbitrary location in expression space (cf. Fig. 1). Instead, consider a bias in the expression of the key that depends on the expression of the lock: *k* = *l* + *f* (*l*), where *f* is an arbitrary function (Fig. 4A). This bias introduces asymmetries in the network connections (Fig. 4B) which cause the activity bump to move (Fig. 4C). The direction of this movement depends on whether the bias *f* (*l*) is positive or negative. In fact, a mathematical analysis [27] shows that the network dynamics can be approximated by a moving bump at a time-dependent position *x*(*t*) in expression space, which changes according to the differential equation d*x/*d*t* = *f* (*x*) (Fig. 4D). Hence, arbitrary dynamics can be innately encoded by a verbatim conversion of differential equations into biases in molecular expression levels.

**Fig. 4:**
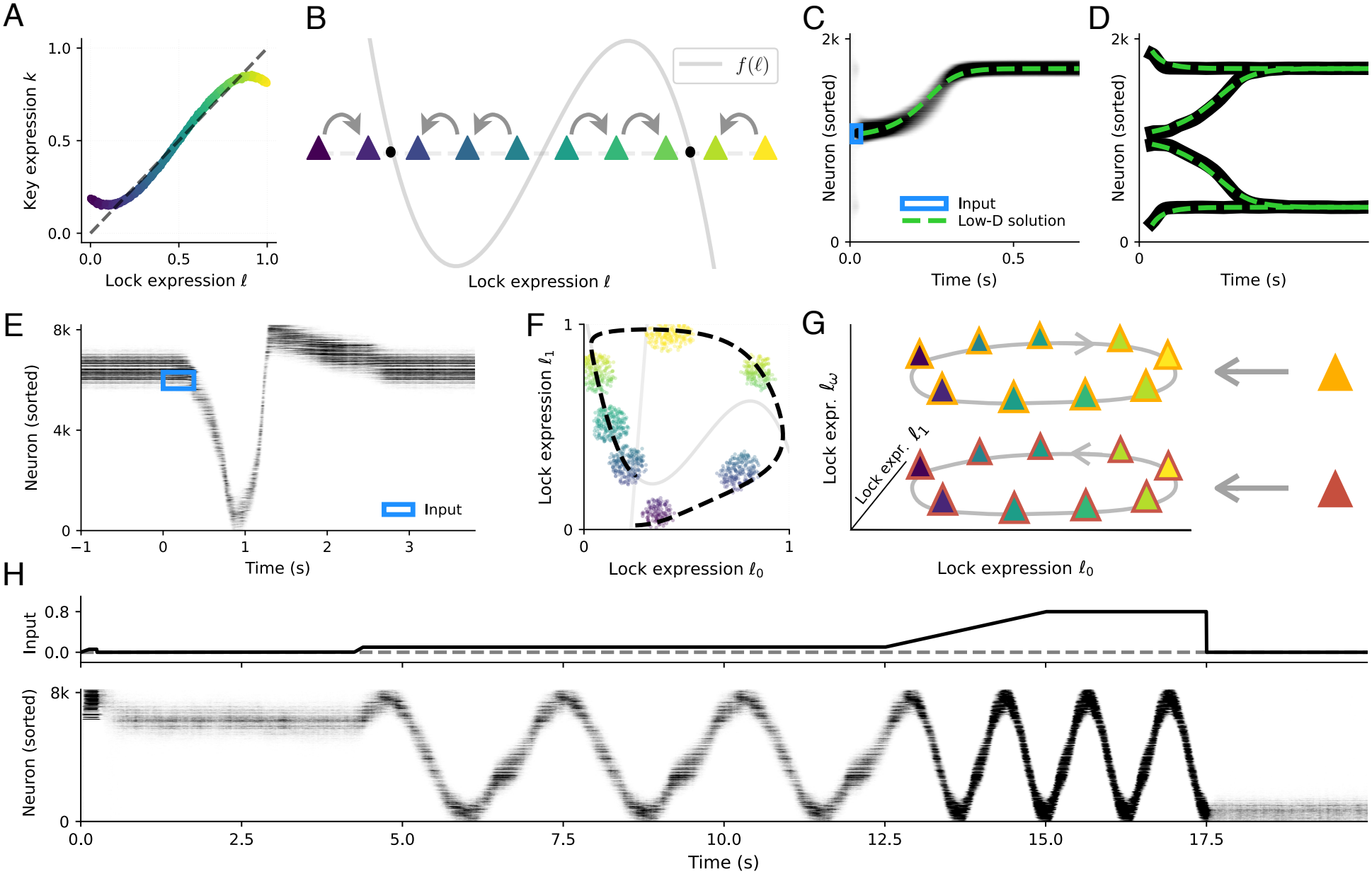
Innate motor programs. (**A**) Continuous expression levels for a single pair of lock and key. key and lock expression are correlated, with an additional bias that depends on the lock expression level: *k* = *l* + *f* (*l*). (**B**) The bias *f* (*l*) (gray line) in key expression gives rise to asymmetries in the connection profile which cause an activity bump to move in a direction (arrows) that depends on the sign of the bias. (**C**) Firing rates of neurons sorted by lock expression in response to a brief input pulse (blue square; centered at *l* = 0.52). The dashed line indicates the solution to the 1D system d*x/*d*t* = *f* (*x*). The small activity bumps close to initialization are due to the two stable attractors. (**D**) Same as (**C**), but for four different initial conditions. (**E-F**) Excitable dynamics: FitzHugh-Nagumo model [46] implemented with two key-lock pairs. (**E**) Dynamics of the network over time. Before stimulation (*t <* 0 s), the network activity sorted by *l*_0_ displays a bump at a stable position. After the network receives a tuned stimulus pulse (blue), the bump follows a stereotypical trajectory and returns to its rest state. (**F**) Snapshots of the activity bump trajectory in the molecular space defined by (*l*_0_, *l*_1_). The black dotted line indicates the trajectory of the 2D dynamical system, light gray lines its nullclines. (**G-H**) Controlled oscillation network. (**G**) Two populations of neurons express two key-lock pairs (*l*_0_, *l*_1_) that generate bump oscillations in opposite directions. The expression of a third, binary lock (*l*_*ω*_) ties these two populations to an input that controls the oscillation frequency. (**H**) Network activity sorted by expression across time (bottom) in response to input (top; difference between the two input populations). In absence of an input difference, the network is a line attractor (different steady bump locations at the beginning and end).

A characteristic of innate behavior is the execution of fixed motor programs in response to a stimulus [1]. To illustrate that such stereotypical responses can be developed by chemoaffinity, we considered a network with two key-lock pairs, in which the expression of the keys is biased by both locks. The structure of the bias is chosen to reflect an excitable dynamical system [FitzHugh-Nagumo [46, 47]]. The network receives sensory input that is organized by the same key-lock system.

After development, the network displays a bump of activity in expression space, the position of which is stable in the absence of sensory inputs (rest state, Fig. 4E). Yet, a transient stimulus can initiate a stereotypical trajectory of the activity bump that lasts several seconds before the network returns to its rest state (Fig. 4E-F, movies S2, S3). This response is highly selective to the stimulus, which must activate neurons in an appropriate region in expression space (movie S4).

Many innate motor programs, such as locomotion, exhibit oscillations with adjustable frequency. Networks with such dynamics can be formed by combining the stochastic expression motifs we introduced earlier. Consider two populations of neurons with biased expression levels such that they generate a clockwise and counter-clockwise oscillation in expression space, respectively (Fig. 4G). The two populations are defined by an additional binary lock that also organizes the input, as in the path-integration system. The network displays oscillations, the frequency of which can be smoothly controlled by the input (Fig. 4H).

Importantly, in both the excitable and the oscillatory system, the phase at which a neuron is active is entirely determined by its expression profile, such that a mapping to motor effectors could also be achieved by molecular labels. In summary, these results suggest that chemoaffinity has the potential to establish innate motor programs by shaping networks with arbitrary temporal dynamics that can be triggered by specific sensory stimuli.

## Discussion

We have shown how unconnected neurons with cell-autonomous, stochastic molecular expression patterns can develop into functional neural networks by means of a simple chemoaffinity rule. The computation performed by the resulting network is determined by the expression statistics. Cognitive functions and motor programs could therefore be innately formed by stochastic cellular differentiation programs.

The expression statistics provide molecular fingerprints for cognitive computations that could serve as a starting point for the identification of candidate molecules. Moreover, the model predicts that the relevant molecular expression levels are directly linked to the semantics of neural representations. If the grid cell attractor [39] was indeed established by chemoaffinity, the relative spatial phase of pairs of grid cells should correlate with differences in molecular expression levels. Motor neurons that are active at the same phase of an innate motor program should share a similar expression pattern.

The number of molecules required even for relatively advanced computations is surprisingly small. This could have a major evolutionary advantage because it may have enabled an efficient selection of functional network architectures in relatively low-dimensional search spaces of molecular systems. As a corollary, it may enable the technical application of evolutionary algorithms to identify neural architectures for cognitive or motor skills, not only for neuroscience but also for robotics and artificial intelligence [48].

While chemoaffinity could be a powerful means for the innate initialization of neural circuits [19], it is likely complemented by activity-dependent mechanisms [16]. Neuronal plasticity—driven by experience [49] or internally generated activity [15]—could not only further refine the network but also bind the activity patterns of the network to sensory data and motor commands [43, 50].

For conceptual clarity, and because we prefer to remain agnostic about which of the known guidance molecules could be at play, we used a simple phenomenological model of chemoaffinity [20]. While it encapsulates the hypothesis that neurons forge connections based on molecular similarity—arising, e.g., from similar birth date [51] or clonal origin [52]—, the model abstracts away much of the molecular and biophysical complexity of guidance systems [53–55]. For example, the phenomenon of matching key and lock expression during the development of topographic maps is mediated by a complex interplay of attractive and repulsive forces [53, 56]. Whether and how these mechanisms operate in local, recurrent networks and which means of developmental self-organization they offer remains to be resolved.

The example systems we presented highlight a peculiar correspondence between the statistical structure of expression levels and the associated neural computation. Discrete expression patterns generate an attractor network that categorized its inputs into discrete classes. Hierarchically clustered expression levels form hierarchies of categories. Random expression levels along a continuum establish continuous attractors, i.e., representations of continuous variables. All these neural computations—categorization, hierarchies, continuous variables—implement structured priors [43] over the statistical or topological nature of the variables that the network encodes. It is intriguing to speculate that the expression of guidance molecules may have evolved to form a statistical mirror image of the animal’s environment and its dynamics.

## Acknowledgments

HS would like to thank Andreas V. M. Herz and Edvard I. Moser for a discussion that sparked this work, and we thank Susanne Schreiber, Johannes J. Letzkus, Joram M. Keijser, Omri Barak, and the members of the Sprekeler Lab for feedback on the manuscript.

## Funding

This work was supported by the Deutsche Forschungsgemeinschaft (DFG, German Research Foundation) under Germany’s Excellence Strategy – EXC 2002/2 “Science of Intelligence”, project number 390523135, and the Collaborative Research Center TR 384 “IN-CODE”, project number 514483642.

## Author contributions

H.S. initiated, conceptualized and supervised the study. F.S. and S.C. performed the numerical simulations and analyzed the simulated data. F.S, S.C. and H.S. wrote the manuscript.

## Declaration of interest

The authors declare no competing interests.

## Data and materials availability

No datasets were generated or analyzed during the current study. All scripts used for simulation, analysis and generation of figures were written in Python. All code will be made publicly available upon publication.

## Supplementary Methods

### Network dynamics

We simulated networks of *N* rate neurons with *N*_*E*_ excitatory and *N*_*I*_ = *N* − *N*_*E*_ inhibitory neurons. Their internal states *x*_*i*_ are determined by the stochastic differential equation

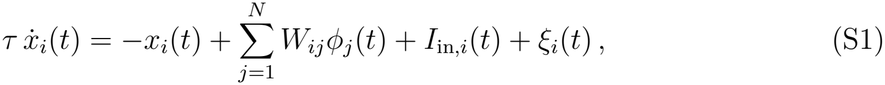

where *W*_*ij*_ is the recurrent synaptic weight from neuron *j* to neuron *i*, and *τ* is the neuronal time constant. The rates are obtained by applying a nonlinearity to the states, here a rectified linear function, *ϕ*_*i*_(*t*) = [*x*_*i*_(*t*)]_+_. In addition to the recurrent input, each neuron is driven by an external input that consists of a constant term *I*_0_ and input neuron rates *ϕ*_in,*j*_(*t*) transmitted via input weights *W*_in,*ij*_:

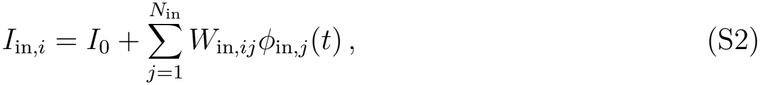

where *N*_in_ is the number of input neurons. We assumed all inputs to be excitatory: *I*_0_, *W*_in,*ij*_, *ϕ*_in,*j*_(*t*) *>* 0. The specific inputs for each model are described below. Finally, every neuron receives Gaussian noise, 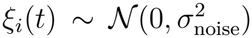, which is uncorrelated in time and between neurons: 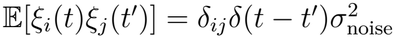.

To simulate the RNN dynamics numerically, we used the Euler-Maruyama method [57] with time step Δ*t*. We chose Δ*t* to be sufficiently small for stable evolution. For a simulation duration *T*, we saved the activity with a recording time step Δ*t*_rec_. The initial states were drawn from a standard normal distribution, *x*_*i*_(*t* = 0) ∼ *N* (0, 1). All simulations were implemented in NumPy [58] and PyTorch [59]. The numerical values for all simulation parameters can be found in Table S1.

### Model of chemoaffinity

We modeled the developmental process of forming connections using an abstract version of Sperry’s chemical affinity theory [20]: Every neuron expresses keys and locks, and the connection between two neurons depends on the match between the expression levels of keys and locks.

### Molecular expression levels determine connectivity

Neuron *i* in our model is characterized by the expression levels of *M* key-lock pairs, 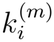 and 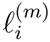 for *m* ∈ {1, …, *M*}. In our models, key and lock expression levels were generally highly correlated. The specific correlation structure for each model is of key importance and is described below.

A synapse from neuron *j* to *i* forms with probability *p*_*ij*_, which depends on the distance in molecular expression space,

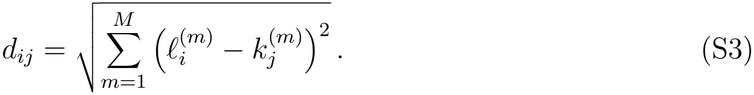

We chose a Gaussian kernel for the connection probability:

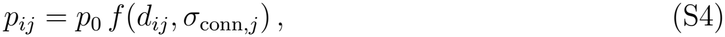

with 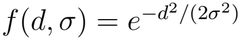. The factor *p*_0_ controls the maximal connection probability, which we increased over time from *p*_0_ = 0 to *p*_0_ = 1 to model the developmental growth process (see below). The tolerance *σ*_conn,*j*_ determines up to which distance in molecular expression space the neuron *j* forms synapses. The tolerance depends on whether the presynaptic neuron is excitatory or inhibitory. Ordering the neurons by type, we defined the excitatory neuron tolerance *σ*_conn,*j*_ = *σ*_conn,*E*_ for *j* ≤ *N*_*E*_, and the inhibitory tolerance *σ*_conn,*j*_ = *σ*_conn,*I*_ for *j > N*_*E*_. Note that in the main text, we use the symbol *σ* for the tolerance, without the subscript “conn” and the specification of the neuron type (which is given by the c ontext). We chose this slight inconsistency to keep the main text simple while allowing us to distinguish *σ*_conn_ from other similar parameters in the Methods.

We defined the recurrent weight matrix as the binary synaptic connectivity matrix *B*_*ij*_ multiplied by weights *w*_*ij*_ :

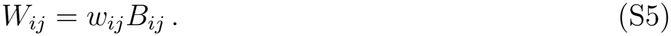

The synaptic connections *B*_*ij*_ are binomial variables, *B*_*ij*_ ∼ *B*(*K, p*_*ij*_), where *K* is the maximum number of synapses between neurons. We used *K* = 1 for all simulations, except for the large-scale 2D path-integration model (Fig. 2F-I). Choosing *K >* 1 reduces the variability in the weight matrix, leading to a more stable pattern for a given network size. We set all excitatory weights to *w*_*ij*_ = *w*_*E*_ *>* 0 for *j* ≤ *N*_*E*_ and inhibitory weights to *w*_*ij*_ = *w*_*I*_ *<* 0 for *j > N*_*E*_.

Input weights are set via the same mechanism, *W*_in,*ij*_ = *w*_in_*B*_in,*ij*_, matching input cell keys to recurrent cell locks. We assumed excitatory inputs and used tolerance *σ*_conn, in_ = *σ*_conn,*E*_ and global connection probability *p*_in,0_ = *p*_0_. The input weights were set to *w*_in_ = 1*/N*_in_.

### Developmental growth

To model developmental growth, we simulated network dynamics Eq. S1 while gradually increasing the global connection probability from *p*_0_ = 0 to *p*_0_ = 1. For a continuous simulation, we gradually added new synapses at every increase in *p*_0_ (instead of resampling the weights every time) and used the last network state as the initial state for the simulation between weight updates.

More specifically, we chose *S*_dev_ time p oints, *t*_*s*_ ∈ [0, *T*], *s* = 0, 1, …, *S*_dev_, to u pdate *p*_0_. We implemented continuous growth by first d rawing the “final” we ights *W*_*ij*_ (*S*_dev_) for the maximal connection probability *p*_0_. We then iteratively defined a set of m asks *M*_*ij*_(*s*) for every increment *s* with *p*_0_(*s*) = *t*_*s*_*/T* : We started with *M*(0) = 0. Then, at every *s*, we counted the number of existing nonzero mask entries, *n*(*s* − 1) = ∑_*ij*_ *M*_*ij*_ (*s* − 1). The number of newly added connections was Δ*n*(*s*) = *N* ^2^*p*_0_(*s*) *n*(*s* − 1). We chose *n*(*s*) of the nonzero entries in *M*(*s* − 1). The new mask *M*(*s*) was a copy of *M*(*s* − 1) with all additional Δ*n*(*s*) chosen synapses set to 1. During the simulation, we updated the weight matrix to *W*_*ij*_ (*s*) = *M*_*ij*_ (*s*)*W*_*ij*_ (*S*_dev_). The last state vector with entries *x*_*i*_(*t*_*s*_ − Δ*t*) was used as the new initial condition, and we ran the dynamics for the interval from *t*_*s*_ to *t*_*s*+1_ until the next update.

### Relation of chemoaffinity model to molecular guidance systems

Our model of the developmental connectivity rule is an abstract, conceptual version of Sperry’s chemoaffinity theory [20]. Sperry’s theory, and hence our model, is of course highly simplified, and many studies have revealed different and more complex molecular mechanisms [22]. Here, we briefly discuss how our model relates to these results and to biophysical models of chemoaffinity, in particular axon guidance.

A well-studied example close to our abstract model is the retinotopic mapping in Fig. 1B. Relevant axon guidance molecules have been identified as Eph receptors (keys) and ephrin ligands (locks) [26]. Taken out of the retinotopic context, these key-lock pairs usually lead to a “binary” interaction mode, in which their presence dictates the connectivity rather than their concentration [60]. Instead, our model captures a “proportional” interaction [60], where differences in expression levels dictate connectivity. In the retinotopic context, this arises as an effective interaction resulting from competition between axons and various other molecular mechanisms [56]. How our simplified chemoaffinity model (uniform distributions, Euclidean distance, Gaussian connection probability) relates to more detailed biophysical models of known guidance systems remains to be explored.

For retinotopic mapping, both Eph and ephrin are expressed in the retina and superior colliculus. Their interaction is repulsive, but their expression is anticorrelated within each neuron. This allows us to map between the biological system and our model by defining *k* = *c*_Eph_ in the retina and *l* = 1 − *c*_ephrin_ in the superior colliculus. The phenomenological expression levels *c*_Eph_ and *c*_ephrin_ may have a nonlinear—e.g., logarithmic—relation to suitably normalized molecular concentrations.

## Model specifics

### Bump attractor

For the bump attractor, we set up a network of excitatory and inhibitory neurons with one key-lock pair (*M* = 1). locks and keys are highly correlated. We drew a joint uniform variable *z*_*i*_ ∼ *U* (0, 1) and set

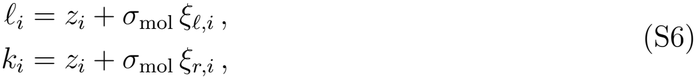

where *σ*_mol_ denotes the spread. Here, and in all similar expressions, *ξ*s are independent standard normal variables.

The distance in expression space, Eq. S3, simplifies to *d*_*ij*_ = |*l*_*i*_ −*k*_*j*_|. Synaptic connections and weights are drawn according to Eqs. S4, S5, with local excitation, *σ*_conn,*E*_ ≪ 1, and global inhibition, *σ*_conn,*I*_ = ∞.

The *N*_in_ input neurons tile the expression space [0, 1] with their keys: 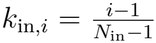. The input neuron rates *ϕ* (*t*_in,*i*_) represent an input pulse in expression space with center *µ*_pulse_ and width *σ*_pulse_. We set the pulse shape to

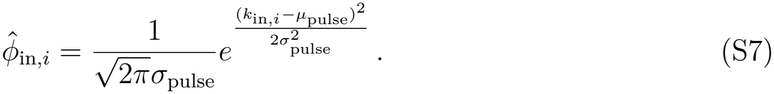

We drew a random input location *µ*_pulse_ ∼ *U* (0, 1) independently for each pulse. We further drew times *t*_pulse_ with pulse duration Δ*t*_pulse_ and set the input rates according to

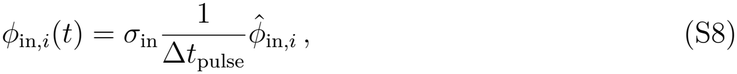

for *t* ∈ [*t*_pulse_, *t*_pulse_ + Δ*t*_pulse_], and *ϕ*_in,*i*_(*t*) = 0 else. We chose parameters *σ*_pulse_ = 0.05, Δ*t*_pulse_ = *τ*, and *σ*_in_ = 1.

For the simulated developmental growth in Fig. 1F-G, we updated the connectivity *S*_dev_ = 101 times and simulated network activity for a period of length Δ*T* = 100 *τ* between each update. The entire simulation thus covered the duration *T* = *S*_dev_Δ*T*. For the input pulses, we chose time points that were not aligned with the weight updates. We partitioned the interval [0, *T*] into 21 blocks, shifted the blocks by half a block width, and added some noise (with amplitude 0.1× block width).

### Path-integration model, 1D

For the one-dimensional path-integration model, we used *M* = 2 different key-lock pairs. Excitatory cells express a discrete head-direction (HD) lock,

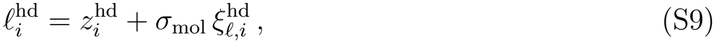

with binary 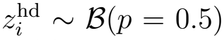. This lock determines the input to the network: “leftward” HD input for 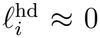, “rightward” HD input for 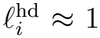 (see below). Excitatory neurons additionally express a “position” lock,

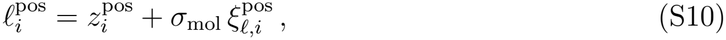

with uniformly distributed 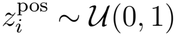. The corresponding key is defined as a noisy copy plus a bias induced by the HD lock expression:

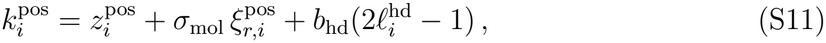

with connectivity bias parameter *b*_hd_ = 0.02. With this connectivity bias, cells that receive input from “leftward” HD cells 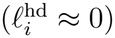 also receive more recurrent input from cells situated to the left (smaller expression) in position key space. Inhibitory neurons express only the position key-lock pair, for which expression levels are drawn as for excitatory cells, but without the bias in Eq. S11.

Connections between all cells are drawn according to their distance in molecular space, Eq. S3. Note that because cells only express a complete pair of key and lock for the “position” type, this distance is given by 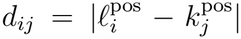, whereas the HD lock (for excitatory neurons) is missing the relevant key for local network connections. For the pathintegration model, we assumed short range excitation and long-range (but finite) inhibition: 0 *< σ*_conn,*E*_ *< σ*_conn,*I*_ *<* ∞. Synapses and weights were again drawn according to Eqs. S4, S5.

Excitatory cells receive input from *N*_in_ head-direction cells, whose identity is determined by the expression level of an HD key, 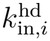. We drew the expression levels from a binary distribution plus noise,

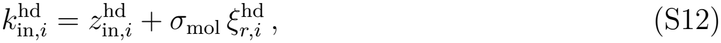

Where 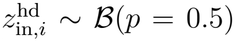. Input cells with 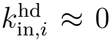 encode the left movement, and those with 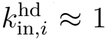 encode the right movement. The input synapses are based on matching the HD key and lock via the distance 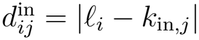 as before, Eqs. S4, S5.

Because we modeled the development before the animal started moving, we assumed that the HD cells fired based on a random input signal. This input signal will later represent the velocity of the animal moving to the left or right on a line (unbounded), an interpretation we used to show the viability of the circuit for path integration. We modeled the input velocities *v*(*t*) as a particle moving in a symmetric double well centered at zero. This approach produces inputs with nonzero speed most of the time, facilitating integration.

The double-well potential is defined as

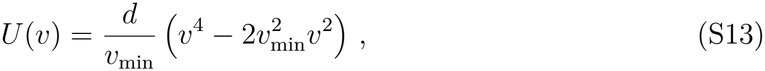

with depth *d* and minima located at ±*v*_min_. The particle dynamics follow an overdamped Langevin equation,

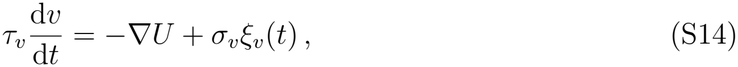

with gradient 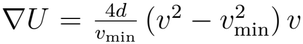 and Gaussian white noise *ξ*_*v*_(*t*). For parameters, we chose depth *d* = 0.5, minima location *v*_min_ = 1, time constant *τ*_*v*_ = 200*τ*, and noise strength *σ*_*v*_ = 1.2. We discretized the stochastic differential equation using the Euler-Maruyama method as for the network dynamics. The initial positions were randomly assigned to one of the two minima. Input neuron rates are defined based on the velocity *v*(*t*) and the HD keys as

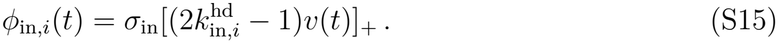

In other words, input cells with HD key 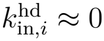 encode negative (“leftward”) velocity, and those with 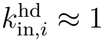 encode positive (“rightward”) velocity.

For the simulated developmental growth in Fig. 2D, we updated the connectivity *S*_dev_ = 21 times and simulated network activity for Δ*T* = 100 *τ* time steps between each update, yielding *T* = *S*_dev_Δ*T* simulation time steps. As initial conditions for simulations between updates, we used the last network state, and for the input, we drew the input for all times *t* ∈ [0, *T*] from one continued process, as described above. Simulating dynamics as if the network was continuously operating throughout development is done for visualization—we do not assume that the time scales for synaptic growth and neuronal dynamics in real development match.

### Path-integration model, 2D

For the two-dimensional path-integration model, we equipped the cells with two HD locks and two key-lock pairs for the position. To limit the network size to be simulated, we set the position locks and keys to grid points (plus noise). We also introduced a slight asymmetry in expression space by drawing one (the position lock in *y*-direction) from a slightly smaller interval. This stabilized the grid pattern along one direction (no symmetry of 90^*°*^ rotation). In biology, we would expect asymmetries between different locks to be the rule, and full symmetry to be the exception (requiring fine-tuning).

For excitatory neurons, we generated a 64 × 56 equally spaced grid of joint variables 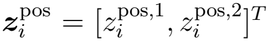 on the rectangle 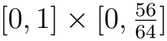. Bold letters indicate 2D vectors, and subscript *i* is the neuron index. We created a copy of this grid for each HD lock combination. For the two HD species, there are four combinations: (0, 0), (0, 1), (1, 0), (1, 1). For each copy, we set the joint HD variables to the relevant combination, e.g., 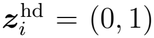. Neurons are indexed according to grid point and copy, so that indices run from *i* = 1 to *i* = *N*_*E*_, with *N*_*E*_ = 64 56 4 excitatory neurons. keys and locks are noisy copies of the joint variables, with a bias depending only on the particular HD lock species. That is,

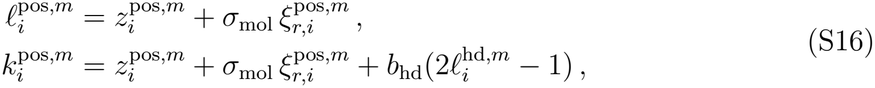

for *m* ∈ {1, 2}, with bias parameter *b*_hd_ = 0.005.

For inhibitory cells, we used a similar grid, but with only 32 × 28 grid points (larger spacing). Although inhibitory cells do not express HD locks, we still used four copies, resulting in *N*_*I*_ = *N*_*E*_*/*4. locks and keys are drawn as for the excitatory cells, but without the bias term in Eq. S16.

The distance for synaptic connections is now the Euclidean norm over both position pairs, 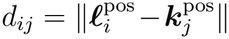. As in the 1D model, HD pairs are not present because local neurons only express HD locks but not keys. The tolerance *σ*_conn_ for both *E* and *I* neurons is assumed to be finite.

The input for the two-dimensional model was generated in a manner similar to the onedimensional model. We used a rotationally symmetric version of the double-well potential (i.e., a ring) by replacing *v* with the norm of the 2D velocity ***v*** in Eq. S13:

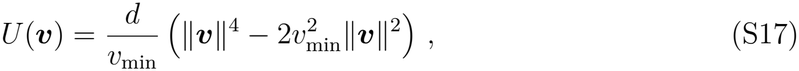

so that the gradient becomes 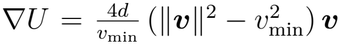. For parameters, we chose depth *d* = 2, minima location *v*_min_ = 1, time constant *τ*_*v*_ = 50*τ*, and noise strength *σ*_*v*_ = 1.2. In 2D, the HD input is given by 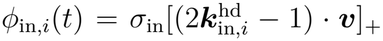. HD cells with four different HD key combinations split the input space into four quadrants. For example, cells with 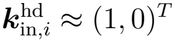 encode velocity in the lower-right quadrant (*v*_1_ *>* 0, *v*_2_ *<* 0), cf. Fig. 2H.

### Autocorrelation analysis (1D and 2D path-integration models)

We investigated whether the innately modeled network was able to perform path integration even before the animal started moving. Thus, we took the random HD input as a “tobe” velocity signal and integrated it to obtain a “to-be” external position, ***p***(*t*). We then computed the autocorrelation of each neuron in the external space: How is the firing rate *ϕ*_*i*_(*t*) at position *p*(*t*) related to the later rate *ϕ*_*i*_(*t*^*′*^) at position *p*(*t*^*′*^)? Specifically, we defined distance bins of width Δ*d* and computed the correlation for all pairs of positions within such a bin,

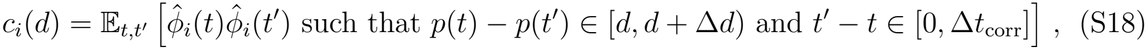

where 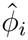 is the z-scored firing rate and 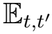 is the empirical expectation over pairs of time points. The parameter Δ*t*_corr_ restricts the average to time points in the near future. This was performed to reduce the influence of accumulating integration errors.

For the 1D model, Fig. 2E, we used distances *d* ∈ [−170, 170], bin width Δ*d* = 0.85 and Δ*t*_corr_*/τ* = 200. For the 2D model, the expressions above are generalized to two separate distances (*d*_1_, *d*_2_). For Fig. 2G, we used *d*_*i*_ ∈ [−140, 140] for *i* = 1, 2, bin width Δ*d* = 8.2, and Δ*t*_corr_ = 500. Because both the 1D and 2D grid cell models have absorbing (non-periodic) boundaries, we only observed a grid-like pattern away from the boundaries. Thus, we restricted our analysis to cells whose locks were at least Δ*l* = 0.15 away from the boundary (cf. Fig. 2F).

### Analysis of toroidal structure in 2D path-integration model

The moving grid-like patterns suggest that the population activity has a toroidal topology similar to that observed by Gardner et al. [39]. To show this, we analyzed our data closely following their steps and using their published code.

We were interested in the toroidal activity structure, which is independent of the current input. Because excitatory neurons are tuned to HD input (cf. Fig. 2H), we restricted the analysis to cells with one type of HD lock, namely 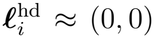, and normalized the rate vector ***ϕ***(*t*) = [*ϕ*_1_(*t*), …, *ϕ*_*n*_(*t*)] with *n* = *N*_*E*_*/*4 at each time point by dividing each entry *ϕ*_*i*_(*t*) by the norm across neurons ∥***ϕ***(*t*)∥. These two steps reduced the influence of the HD input and allowed a clearer observation of the toroidal structure.

We applied principal component analysis (PCA) to the normalized rates. An ideal moving hexagonal grid with periodic boundary conditions would create a six-dimensional manifold [39]. Thus, we projected the rates onto the leading six PCs, which contained 39 % of the variance (Fig. S1A). The projected data were downsampled by first selecting the 24,000 time points with the largest L1 norm. In the second step, we applied fuzzy downsampling, a topological downsampling method that removes outliers [39].

The remaining 1024 data points in six dimensions were subjected to persistent cohomology analysis using of the Ripser package [61]. This resulted in persistence diagrams (Fig. S1C-D). Two *H*^1^ complexes (rings) had exceptionally long lifetimes. We defined the angles associated with these as *φ*_1_, *φ*_2_. Coloring the data projected onto the leading 6 PCs by these coordinates shows that there is an alignment between the low-dimensional structure, and the torus angles (Fig. S1E). Coloring the integrated HD input with these angles indicates that they tile this external space in two directions, despite imperfect integration (Fig. S1F). Single neurons show localized tuning to the cohomology angles (Fig. S1G). Finally, we applied the non-linear dimensionality reduction UMAP [62] with cosine metric and parameters *n* neighbors = 100, minimal distance = 0.5. This also shows the twisted toroidal structure [39] and the alignment between coordinates *φ*_1_, *φ*_2_ to the torus angles (Fig. 2I).

### Discrete attractors: evidence integration

We designed the attractor networks in Fig. 3A-G by assigning highly correlated key-lock pairs with binary expression levels (plus noise). For each pair *m* ∈ [1, …, *M*], we drew a joint variable 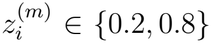 with equal probabilities. lock and key expression levels are noisy copies of this joint variable,

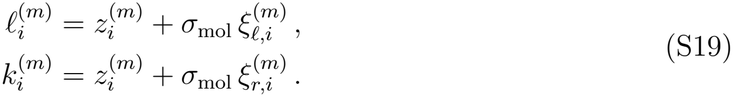

As before, weights depend on the Euclidean distance in expression space, Eq. S3, and a Gaussian kernel for the connection probability (Eqs. S4, S5). We set a small excitatory neuron tolerance, *σ*_conn,*E*_ ≪ 1, and infinite inhibitory neuron tolerance *σ*_conn,*I*_ = ∞, resulting in global inhibition and two assemblies with internal recurrent excitation.

For the evidence integration model, Fig. 3E-F, we used *M* = 1 key-lock pairs. Each neuron receives excitatory input *I*_in,*i*_(*t*) = *I*_0_ + *I*_*i*_(*t*), which is given by the sum of a constant term and a time-varying signal. The time-varying input *I*_*i*_(*t*) to a neuron uniquely depends on its lock expression level. Specifically, we implemented the inputs by two input neurons with low and high key expression, respectively. Input weights are set so that the input neuron with high key expression projects exclusively to excitatory neurons with high lock expression level *l*, and vice versa. As a result, excitatory neurons with similar lock expression receive identical external inputs, which are chosen as a stochastic signal with mean *I*^*l*^, variance *D*_*I*_ and duration of Δ*t* = 3 s: 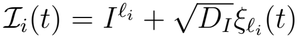 where *ξ*_*l*_ (*t*) is a white noise process. To test the stability of the final activity patterns, we simulated the network for an additional 1.5 s after the varying inputs were switched off. We computed the choice probability by providing only white noise as input to the network (*I*^*l*^ = 0) and repeating the integration for *n* = 5000 independent trials.

To measure how the network complexity scales with the number of guidance molecules, Fig. 3G, we simulated networks of *N* neurons with *M* lock-key pairs. We provided the network with a short input pulse (Δ*t*_pulse_ = 1 s) tuned to neurons with lock expression levels around a given value ***µ***_pulse_. The input to each neuron is given by

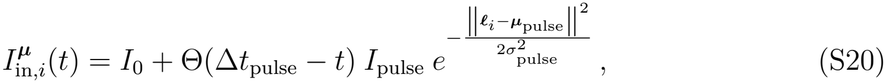

where *σ*_pulse_ denotes the input width. After the input period, we simulated the network activity for additional 2.5 s and measured the average rate 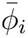 for each neuron in the last 0.4 s. The final configuration is defined by which neurons are active, 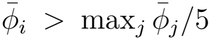, and which are inactive, 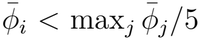. We estimated the number of stable attractors by counting the number of distinct final configurations after sampling the input centers ***µ***_pulse_ from a uniform distribution 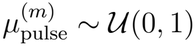.

### Hierarchical model

For the hierarchical model, we considered neurons with *M* pairs of L-R molecules. For each neuron *i* we drew an M-dimensional variable, with components 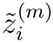 randomly drawn in {−1, 1}. We scaled each axis independently 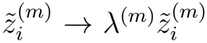. We imprinted a hierarchical structure by choosing the scaling constants *λ*^(1)^ *< λ*^(2)^ *<* · · · *< λ*^(*M*)^ such that the first key-lock pair represents the lowest level in the hierarchy and the last key-lock pair represents the highest level. We obtained joint variables between [0 1] by proper scaling 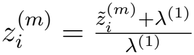.

Given the joint variables 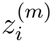, we drew keys and locks as before Eq. S19. We set neural weights by their distance in expression space, Eqs. S3, S4, S5, using finite connectivity width for excitation and global inhibition.

For Fig. 3H-I, we ordered neurons by performing hierarchical clustering on their molecular expression levels (both key and locks) using the unweighted pair group method with an arithmetic mean.

To model development, Fig. 3J, we simulated the network by increasing the global connection probability *p*_0_ from zero to one in 30 steps. For each value of *p*_0_, we provided eight different stimulus pulses *I*^*µ*^(*t*) localized in lock space, Eq. S20. We let the network reach a steady state, simulating for Δ*t* = 3 s after the stimulus terminated. We defined the final network configuration as the average firing rates 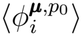 over the last 0.4 s and estimated the number of stable attractors by counting the number of distinct final configurations. We then selected four representative developmental stages and applied multidimensional scaling of the corresponding final network configurations 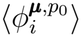.

We read out the category represented by the network activity with a set of output neurons that are connected to the excitatory population via the same guidance mechanism. A subset of output neurons expresses only the first lock and receives input from the excitatory population according only to the expression of this guidance molecule. This subset will therefore code for the higher-level category in the hierarchy. A second subset, instead, expresses the first and second locks, coding for the intermediate level. A final subset of output neurons expresses all three locks and codes for the lower-level category in the hierarchy. In all three subsets, the expression levels of each lock were sampled from the same distributions used in the excitatory population.

### Neuronal sequences

We constructed a model exhibiting neural sequences by equipping neurons with *M* key-lock pairs. The first pair has continuous expression levels *z*_*i*_^(1)^ ∈ *U* (0, 1). For this pair, locks and keys are *biased* copies,

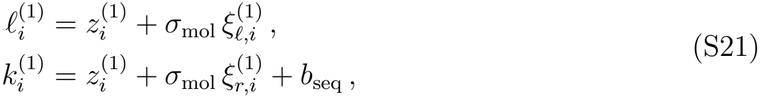

with constant bias *b*_seq_ = 0.03 All remaining *M* − 1 molecular pairs were noisy copies of binary 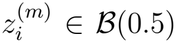, as in Eq. S19. Connectivity is drawn by the distance in expression space containing all *M* species, that is, both continuous and binary.

As shown in Fig. 1, uniform connectivity without bias induces a bump. The additional constant bias *b*_seq_ induces a drift in the bump. As before, for the fixed points, Fig. 3G, the binary key-lock pairs split the network into 2^*M −*1^ subnetworks. Here, each subnetwork can give rise to a sequence. Global inhibition induces a winner-take-all scenario in which only one sequence is active at the time. Finally, when the bump reaches one end of the interval in *l*^(*m*)^, it cannot move further and slowly decays, so that a new bump (and sequence) can arise (selected at random due to the network noise).

### Innate motor programs

To obtain an activity bump, we designed the network connectivity profiles with local excitation and global inhibition with *σ*_conn,*E*_ ≪ 1 and *σ*_conn,*I*_ = ∞. The bump is initialized providing a short input pulse that is localized in lock space, defined in Eq. S20.

The illustrative model in Fig. 4A-D is realized by a network of neurons that express one key-lock pair from a continuous distribution with support [0, 1]. The expression level of the key is correlated to the one of the locks, with a small bias that depends on the lock expression level according to

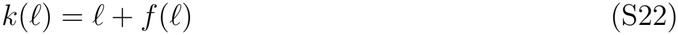

where *f* (*l*) = 1*/τ*_*µ*_ (3*l* − 3*/*2 − (3*l* − 3*/*2)^3^) and *τ*_*µ*_ = 10.

We designed an excitable motor program with a network of neurons that express two continuous L/R pairs:

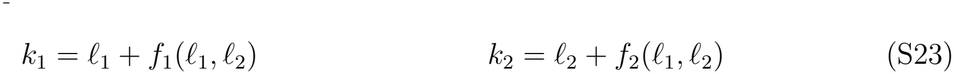

We chose the equations of the FitzHugh-Nagumo (F-N) model as the bias functions:

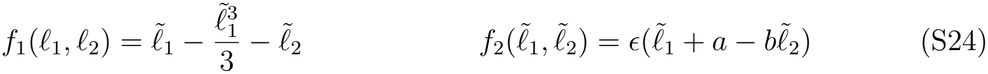

The variables 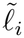 represent the F-N variables after proper scaling and translation in order to have excitable dynamics within lock distribution support [0, 1]. Simulations in Fig. 4E-G are obtained using the following transformation: 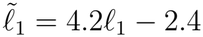 and 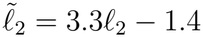.

We modeled controllable bump oscillations by equipping neurons with three key-lock pairs. (*l*_1_, *l*_2_) are continuously expressed in [0, 1] and define the continuous molecular space where the oscillation occurs. The discrete expression of a third lock *l*_*ω*_ determines the structure of the bias functions *f*_1_, *f*_2_, such that neurons with high and low *l*_*ω*_ give rise to clockwise and anti-clockwise oscillations, respectively. Specifically, we chose bias functions that correspond to the real and imaginary parts of a canonical Hopfield oscillator:

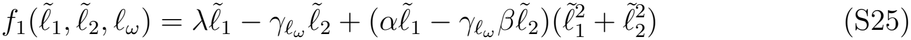

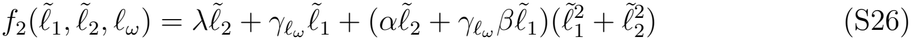

where 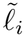 are the expression levels transformed from [0, 1] to [−1.1, 1.1]. Positive parameter *λ* and negative parameter *α* ensure sustained oscillations, with pulsation 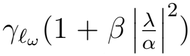.

The two opposite oscillations are given by setting 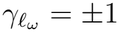 depending on whether *l*_*ω*_ *<* 0.5 or *l*_*ω*_ *>* 0.5. We stimulated the neurons with a time-varying input that depended on the expression level *l*_*ω*_. Neurons with similar *l*_*ω*_ values receive identical inputs. The oscillation frequency is controlled by the difference in the inputs to the neurons with high and low *l*_*ω*_.

## Theory

### Effective radius of weights in expression space

When lock and key expression levels are noisy copies, Eq. S6, then the chemoaffinity mechanism of our model yields local connectivity in expression space, cf. Fig. 1C, H. Here, we showed how the noise between lock and key expression levels and connectivity tolerance gave rise to an effective width of connectivity.

The connectivity probability, Eq. S4, depends on the distance between the postsynaptic lock and presynaptic key expression, Eq. S3. However, to understand the width of connectivity in expression space, we needed a joint basis for pre- and postsynaptic neurons. We chose the lock expression levels ***l***_*i*_, which determine the input to a neuron. The relevant distance for this coordinate system was 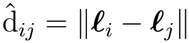.

Because of the noise *σ*_mol_ between keys and locks, the set of presynaptic neurons with locks ***l***^*′*^ will have key expression ***k***^*′*^ distributed around the lock expression. This distribution effectively increased the width of connectivity. The key-lock noise is independent of the tolerance *σ*_conn_, so the corresponding variances add, and we have the effective variance

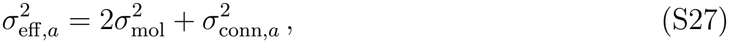

where *a* ∈ {*E, I*} is the neuron type of the pre-synaptic neuron and the factor 2 arises because we defined both *l* and *k* as noisy copies of a joint variable *z*, Eq. S6. Then the connection probability is given by

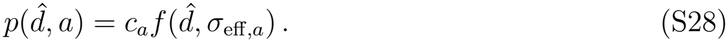

The prefactor 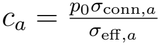 arises from the convolution of the Gaussian distributions.

### Theory for innate motor programs

Here, we show how lock-key-based connectivity leads to a moving bump whose center *µ* moves according to a low-dimensional system *µ*? = *f* (*µ*). For simplicity, we formulate the derivation for a one-dimensional system; however, our results readily extend to multiple dimensions.

We begin with an intuition of how the bump dynamics follow *f* (*µ*). According to our connectivity model, neurons with key *k*^*′*^ send excitatory inputs to neurons with lock *l* = *r*^*′*^ (here and in the derivation below, the prime always indicates presynaptic neurons). If the keys have a slight bias with respect to the locks, *k*(*l*) = *l* + *b*, then a neuron at the bump center, *l*^*′*^ = *µ*, sends excitation to neurons with locks *l* = *k*(*l*^*′*^ = *µ*) = *µ* + *b*. This causes the bump to move in the direction of *b*. If we replace *b* with a function *f* (*l*), then we would expect that the features of the function would be reflected in the dynamics. For example, if *f* (*l*) *>* 0 for *l < l*^*\**^ and *f* (*l*) *<* 0 for *l > l*^*\**^, we would expect the bump to settle at *l*^*\**^. The following derivation makes this intuition precise.

We assume a continuum of uniformly distributed neurons in expression space, *x*_*i*_ *x*(*l*). Weights are correspondingly expressed as *W*_*ij*_ → *w*(*l, l*^*′*^)d*l*^*′*^, and *w*(*l, l*^*′*^)d*l*^*′*^ is the average of the weights from all neurons at lock concentration *l*^*′*^ ∈ [*l*^*′*^, *l*^*′*^ + d*l*^*′*^). Note that the shape of this weight function is determined by both the tolerance of the axon guidance system and the noise in the expression of keys and locks (see the previous section).

The relevant distance in our model is the discrepancy between the postsynaptic locks *l* and presynaptic keys *k*^*′*^ = *k*(*l*^*′*^). Assuming translation invariance in the connection rule, we therefore have *w*(*l, l*^*′*^) = *w*(*l* − *k*(*l*^*′*^)). Note that we assume excitatory and inhibitory activity to be statistically identical, so that we do not need to make excitatory and inhibitory weights explicit.

With the continuum approximation, the dynamics read as

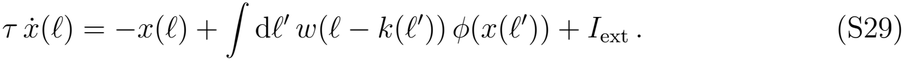

We first express the bump Ψ_0_(*l* − *µ*) to the unperturbed dynamics, that is, perfect unbiased correlation between locks and keys, *k*(*l*^*′*^) = *l*^*′*^. The bump fulfills 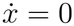, or

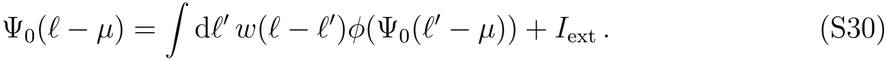

We now introduce the perturbation, *k*(*l*^*′*^) = *l*^*′*^ + *f* (*l*^*′*^). For small *f* of order *O*(*ϵ*), we can approximate the weights as *w*(*l* − *k*(*l*^*′*^)) = *w*(*l* − *l*^*′*^ − *f* (*l*^*′*^)) = *w*(*l* − *l*^*′*^) *∂*_*l*_*w*(*l* − *l*^*′*^)*f* (*l*^*′*^)+*O*(*ϵ*^2^). Inserting this into the dynamics, we have

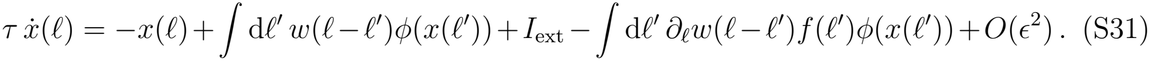

The first three terms sum to zero for *x*(*l, t*) = Ψ_0_(*l* − *µ* (*t*)), assuming that Ψ_0_ i s a solution to the unperturbed dynamics. We are then left with

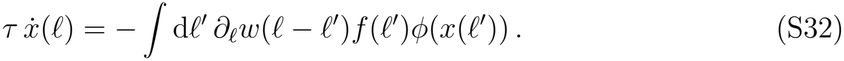

Assuming that the bump is narrow compared to the scale on which *f* changes, we can further approximate the function *f* around the bump center,

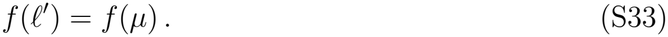

This allows pulling *f* outside of the integral,

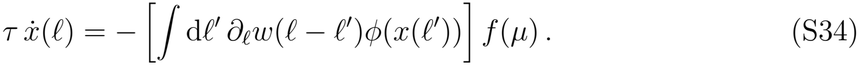

We further express the bump dynamics by the movement of the bump center:

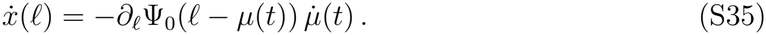

The shape of the bump can be explicitly stated via Eq. S30, (S35)

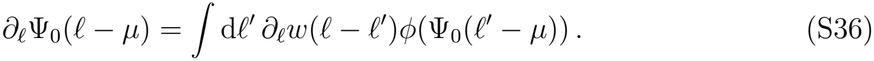

Combining this with Eqs. S34, S35, we finally get

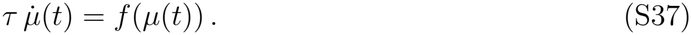

**Fig. S1:**
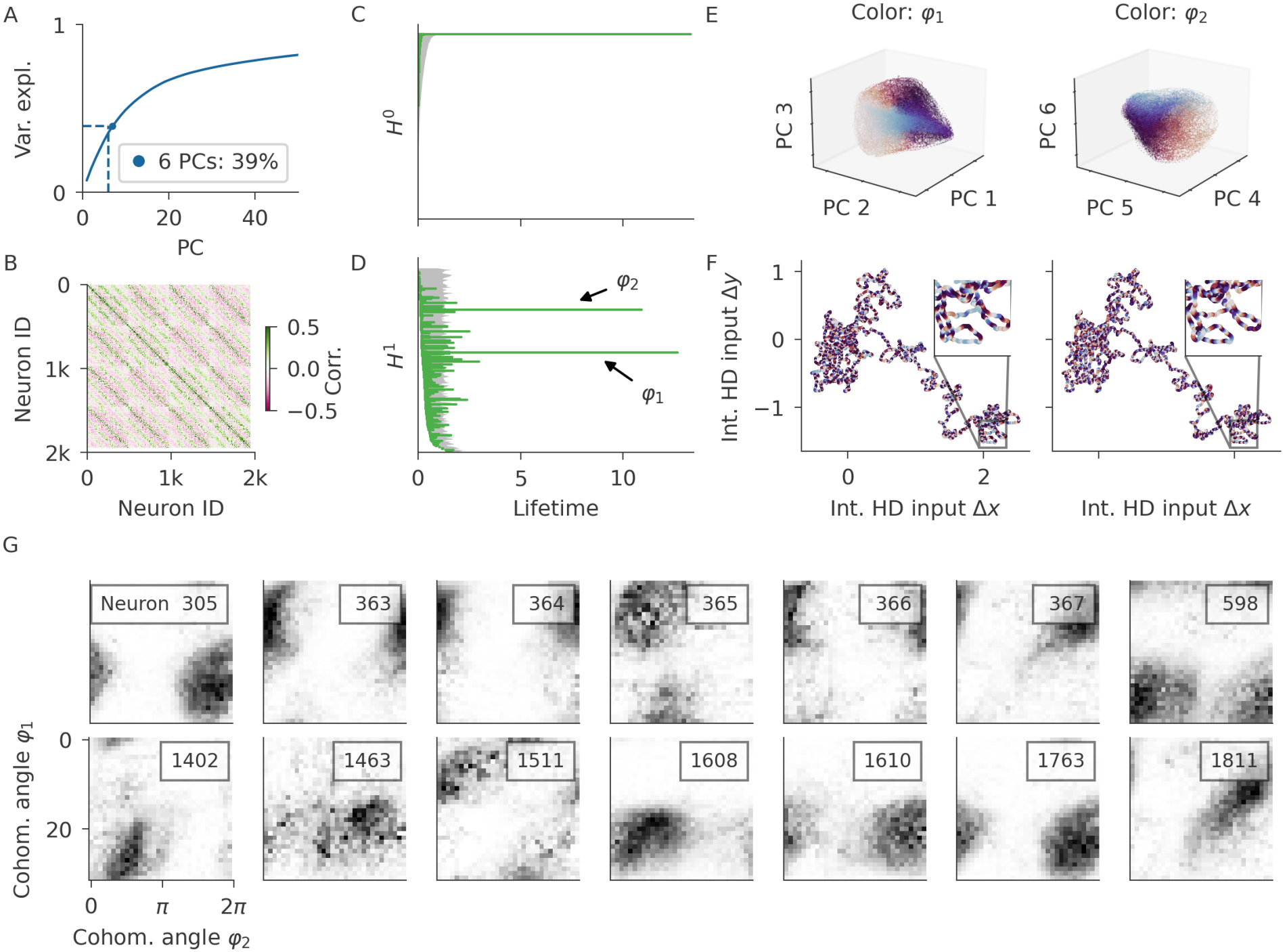
Analysis of toroidal structure in 2D grid cell model. (**A**) Cumulated variance explained ratio of the first 50 principal components (PCs). The first 6 PCs contain 39 % of the variance. (**B**) Correlation matrix of neural activity (excitatory cells with HD lock combination (0, 0) away from boundary). (**C-D**) Results of persistent cohomology analysis: birth and death times of complexes in zero (**C**) and one (**D**) dimension. The two longest-living, annotated *H*^1^ complexes correspond to the two rings defining a torus. Grey shades indicate longest life times for shuffled data (100 independent shuffles, where the rate sequence of each neuron is rolled by an independent random amount of time). (**E**) Projection of the data onto the leading three PCs colored by *φ*_1_ (left) and the next-to leading three PCs colored by *φ*_2_ (right). (**F**) Integrated HD input displays a trajectory in external space (or what will be external space once the animal starts moving). Coloring by *φ*_1_ (left) and *φ*_2_ (right) reveals stripe-like pattern (see insets). Integration is imperfect, as crossings do not always display the same angle. (**G**) Tuning of individual neurons to cohomology angles *φ*_1_, *φ*_2_, measured by firing rate averaged over angle bins (32 bins per angle). Neurons close to each other in expression space (with nearby indices) often, but not always, have similar tuning (neurons 363-366 chosen manually for illustration, all others at random).

**Fig. S2:**
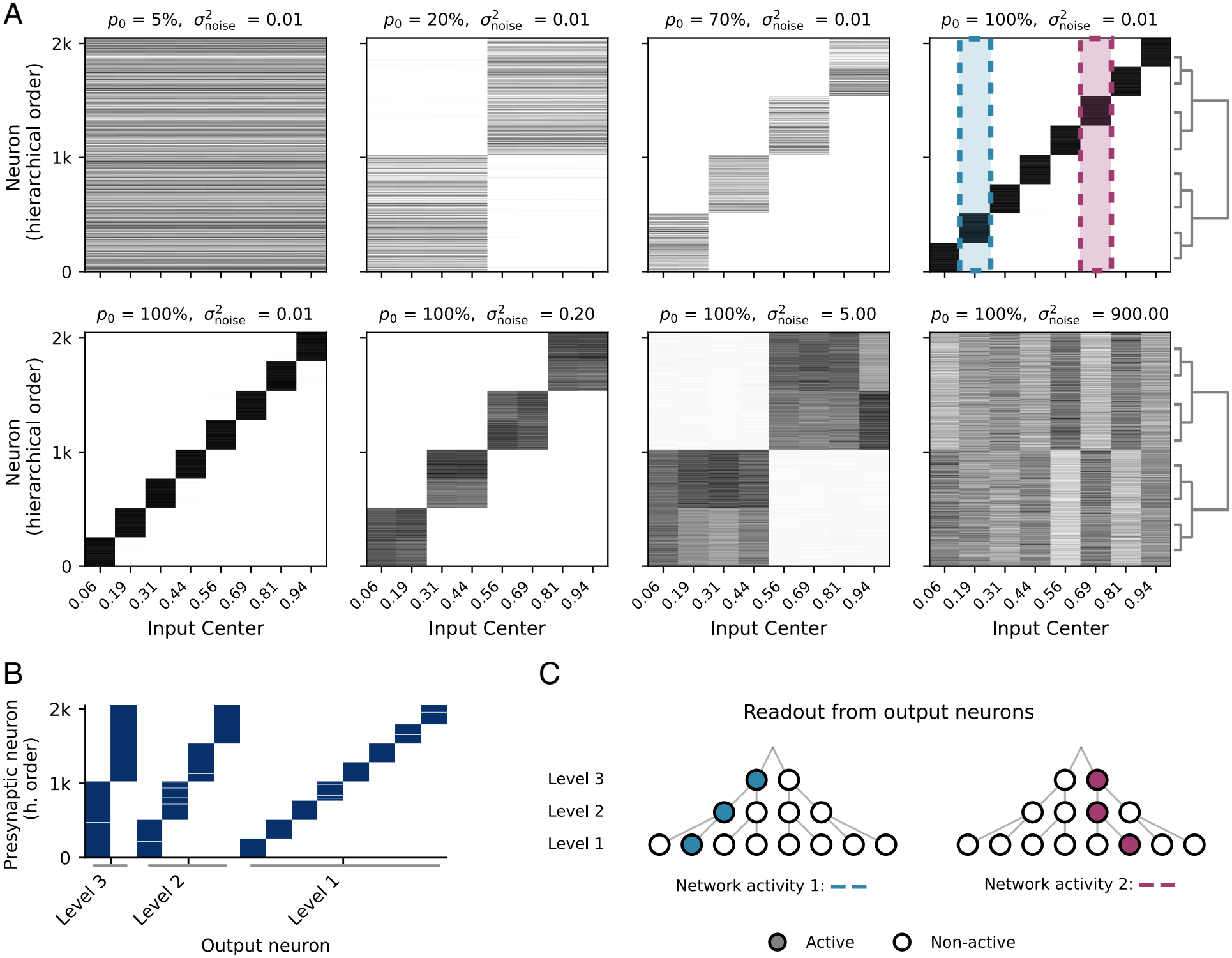
Hierarchical expression of guidance molecules. (**A**) Stable activity configurations after providing stimulus pulses tuned at different input centers in lock space (horizontal axis). (Top row) At initial stages of development, the network reaches a stable activity pattern which is independent of the stimulus tuning (top left). When more synapses are added the stable attractor splits in multiple ones (left to right), according to the hierarchical structure imprinted by the molecular expression, shown by the dendrogram. (Bottom row) At full development, higher-level attractors can be reached by increasing the noise level in the neuronal dynamics (left to right). (**B**) Connectivity matrix between the excitatory populations - sorted by hierarchical clustering - and the output neurons, sorted by their level in the hierarchy. (**C**) Readout of two network activities highlighted in (**A**) by the colored, dotted lines. Output neurons (black circles) are placed in a hierarchical tree according to which hierarchical level they encode (vertical arrangement), and to their molecular expression levels (horizontal arrangement). The activation of output neurons (colored center) allows to decode all levels in the hierarchy.

**Table S1:**
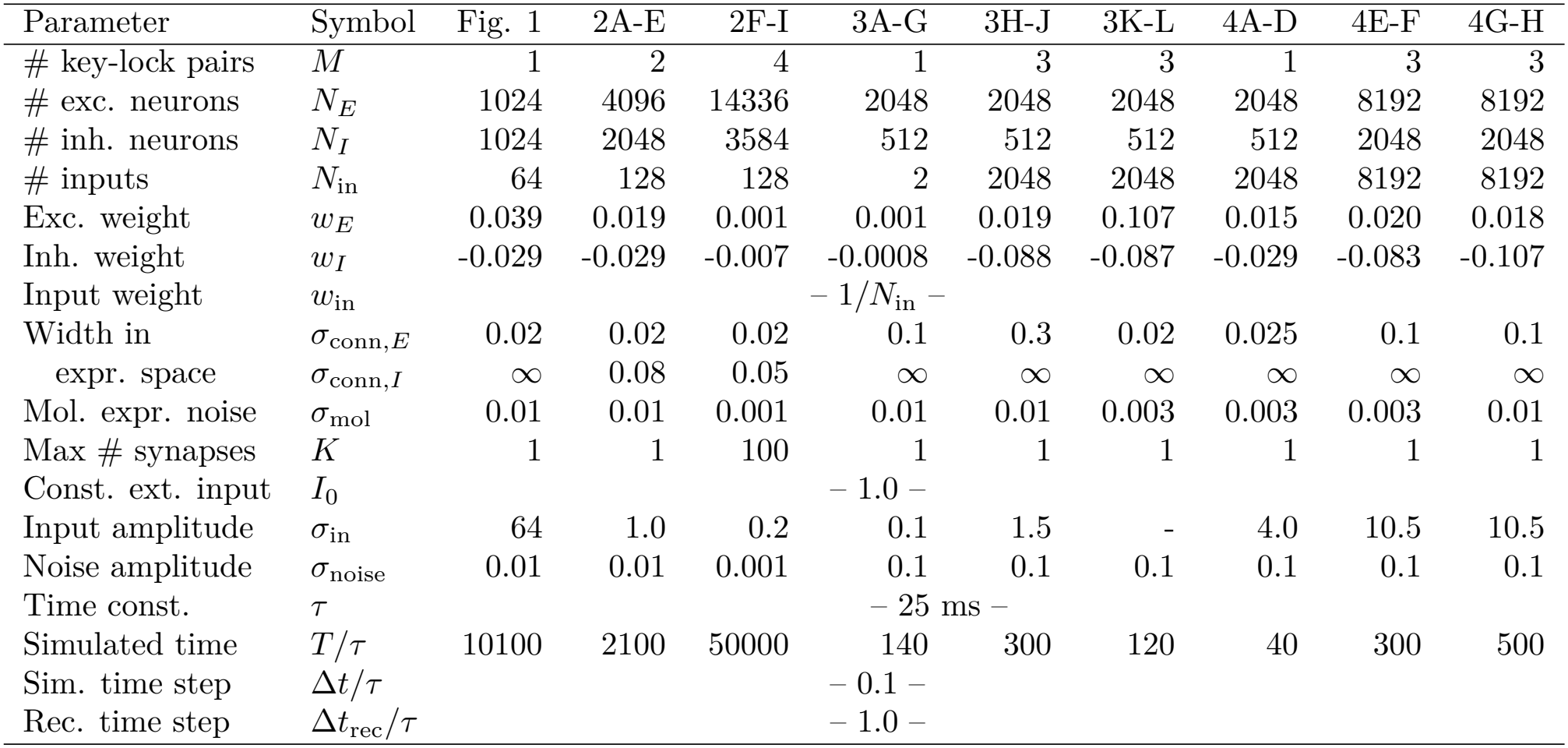
Simulation parameters for all models. Column one and two indicate the variable, all other columns specify the figure c orresponding to the model.

### Caption for Movie S1.

**Neural activity in 2D path-integration model**. Movie of neural activity in the 2D path integration model over time for neurons in physical space (unordered; left) and in molecular space (positioned by lock expression; right). The red bead indicates the integrated HD input (“velocity”) scaled to match the movement of the bumps in molecular space. The trail behind the bead indicates the last 5 s. For our simulation, we provide random input. The close correspondence between grid and integrated input shows that the network readily performs path integration once the input is replaced with actual velocity of the modeled animal. Animated version of Figure 1.

### Caption for Movies S2-S4

**Neural activity in network model with excitable dynamics**. Movie of neural activity (white dots) in a network with excitable dynamics (FitzHugh-Nagumo model). (**S2**) A specific stimulus (stimulated neurons colored in red) elicits a pattern of activity that lasts several seconds, before returning to a rest state. Activity visualized in anatomical space. (**S3**) Sorting the neurons by their expression levels reveals a stable activity bump. After the stimulus presentation, the bump follows a stereotypical trajectory. (**S4**) Stimulating the network with non-specific input does not elicit a response.

## Notes

### Competing Interest Statement

The authors have declared no competing interest.

### Summary of Updates

We revised the terminology (keys and locks vs receptors and ligands) to clarify the abstract nature of our concept. Figure 1 underwent revision, in particular the schematics.

